# The genomic ancestry, landscape genetics, and invasion history of introduced mice in New Zealand

**DOI:** 10.1101/234245

**Authors:** Andrew J. Veale, James C. Russell, Carolyn M. King

**Author notes:** Present address: Department of Zoology, University of Otago, 340 Great King St, Dunedin 9016, New Zealand.

## Abstract

**Summary:** The house mouse (*Mus musculus*) provides a fascinating system for studying both the genomic basis of reproductive isolation, and the patterns of human-mediated dispersal. New Zealand has a complex history of mouse invasions, and the living descendants of these invaders have genetic ancestry from all three subspecies, although most are primarily descended from *M. m. domesticus*. We used the GigaMUGA genotyping array (~135,000 loci) to describe the genomic ancestry of 161 mice, sampled from 34 locations from across New Zealand (and one Australian city - Sydney). Of these, two populations, one in the south of the South Island, and one on Chatham Island, showed complete mitochondrial lineage capture, featuring two different lineages of *M. m. castaneus* mitochondrial DNA but with only *M. m. domesticus* nuclear ancestry detectable. Mice in the northern and southern parts of the North Island had small traces (~2-3%) of *M. m. castaneus* nuclear ancestry, and mice in the upper South Island had ~7-8% *M. m. musculus* nuclear ancestry including some Y-chromosomal ancestry – though no detectable *M. m. musculus* mitochondrial ancestry. This is the most thorough genomic study of introduced populations of house mice yet conducted, and will have relevance to studies of the isolation mechanisms separating subspecies of mice.

## 2. Introduction

The house mouse, *Mus musculus*, provides a powerful model system for understanding evolution, and is arguably the best mammalian model for studies of the genomic basis for reproductive isolation during the early stages of speciation. It includes at least three closely related subspecies with parapatric distributions: *M. m. musculus* found in Eastern Europe and Northern Asia, *M. m. castaneus* in Southeast Asia and India, and *M. m. domesticus,* in western Europe, the Near East, and northern Africa [1]. These three subspecies rapidly diverged in allopatry around 350,000 years ago [2-4], and evidence suggests that *M. m. castaneus* and *M. m. musculus* are more closely related to each other than either is to *M. m. domesticus* [5, 6]. During the past 10,000 years, house mice have become commensal with humans, and as stowaways with them, have become the most successful small mammal colonizers of new continents during the past few hundred years [7, 8].

Regions of secondary contact and introgression may mark where mouse subspecies meet in nature. The best studied of these is a narrow hybrid zone between *M. m. domesticus* and *M. m. musculus* that stretches from Denmark to the Black Sea in central Europe [9-18]. This hybrid zone is young, with mice having colonized this area around 3000 years ago [19, 20]. Hybridization in the wild between *M. m. domesticus* and *M. m. castaneus* is best known from one study of an introduced population in California [21] and one in New Zealand [22]. Within the native range, other possible *domesticus*/*castaneus* hybrid zones in Iran [23-25] and in Indonesia [26] have produced only preliminary results, because these regions are complex, supporting multiple (and potentially undescribed) subspecies [24], and because comprehensive nuclear loci have not been used to look at the levels of admixture across the genomes.

Hybrids between subspecies have been extensively studied in laboratory strains of mice, with data indicating that *M. m. musculus* and *M. m. domesticus* are largely reproductively isolated [27, 28]. These studies have helped us to understand the genetics of speciation, particularly the genetic basis of hybrid male sterility [29-32], revealing an important role of the X chromosome in producing reproductive incompatibilities [33-40]. Studies of both wild and laboratory mice have also found that hybrid male sterility has a complex basis, involving many genes [31, 32, 38, 41]. Laboratory crosses between *M. m. domesticus* and *M. m. musculus* led to the identification of Prdm9 on chromosome 17, the only gene at present known to contribute to hybrid sterility in vertebrates [29, 42]. The identification of other genes underlying hybrid male sterility in the wild remains a challenge, but the combination of mapping studies in the lab and in regions showing limited introgression in nature have identified good candidates for future study [18, 32, 43]. Despite the high degree of hybrid incompatibility and reduced fitness, most standard inbred strains of laboratory mice have been derived from admixtures between mouse subspecies. They often feature Y-chromosome or mitochondrial capture, where these uni-parentally inherited markers do not match the ancestry of the rest of the genome [44-46].

The GigaMUGA array is the third generation of the Mouse Universal Genotyping Array (MUGA) and consists of a 143,259-probe Illumina Infinium II array developed specifically for the house mouse. These probes were designed to be evenly distributed across the 19 autosomal, and X and Y chromosomes with minimal linkage disequilibrium, and they include markers across the mitochondrial genome [46]. While the GigaMUGA array was optimized for Collaborative Cross and Diversity Outbred populations, for substrain-level identification of laboratory mice, SNPs informative for subspecies of origin were also included to facilitate studies of wild mice. The array was designed to have a density of at least one “diagnostic marker” per 300 kb for each subspecies, and to place at least one diagnostic marker for each subspecies within each recombination of the intervals identified in [47]. Therefore, this cheap, high-density array specified for high-throughput biomedical and developmental genomic studies has the potential to analyse colonisation patterns and evolutionary genomics of wild mice at an unprecedented scale.

### Mice in New Zealand

House mice have accompanied humans around the world for thousands of years [48]. Because of their high-standing genetic diversity, it has been possible to track the origins of introduced mouse populations [2], revealing activities and movements of people invisible to traditional historical methods [49-53]. New Zealand was entirely free of all terrestrial mammals until the introduction of the Pacific rat (*Rattus exulans*), which arrived with Polynesian settlement around 1280 AD [54]. House mice arrived among infested food and cargo on early European vessels, starting around the 1790s [55].

Most New Zealand mice closely resemble *M. m. domesticus* morphologically, however some morphological characteristics of *M. m. musculus* have been identified at low frequencies [49]. The mitochondrial diversity of New Zealand mice is surprisingly large, with 23 *M. m. domesticus* D-loop haplotypes descending from all six major *M. m. domesticus* clades, six *M. m. castaneus* D-loop haplotypes from a single clade, and one *M. m. musculus* haplotype so far identified [49, 56]. Across most of the two main islands, and on most offshore islands, *M. m. domesticus* mitochondrial haplotypes predominate. In the southern South Island however, one *M. m. castaneus* mitochondrial DNA is solely found, with a narrow ‘hybrid’ zone around 50 km wide separating the *M. m. castaneus* to the south and *M. m. domesticus* to the north [22, 56]. The same *M. m. castaneus* mitochondrial DNA haplotype is also present in the lower North Island around the Wellington region, and a second one is the only mtDNA haplotype so far identified on Chatham Island. The only place where *M. m. musculus* mitochondrial DNA has been detected is in the lower North Island in Wellington.

While the distribution of mouse mitochondrial lineages across New Zealand has been well documented, the nuclear genomic ancestry of mice in New Zealand is poorly understood. All studies to date have found a predominance of *M. m. domesticus* nuclear ancestry across the country, including ‘hybrid’ populations containing unquantified mixes of the other two subspecies present but insufficiently characterised. Of the few nuclear markers that have been sequenced previously, all mice regardless of mitochondrial haplotype, have had predominantly *M. m. domesticus* ancestry, though some mice in the upper South Island also have also had *M. m. musculus* markers [22, 49]. No *M. m. castaneus* nuclear ancestry has yet been detected in New Zealand mice.

New Zealand mouse populations are of particular interest for genetic studies due to the presence of hybrids between all three subspecies. Hybrids of *domesticus*/*castaneus* are of particular interest, as this mixture has rarely been studied, and has never been confirmed in their native range. In this study, we aimed to ascertain the relative contribution of each subspecies to these hybrid populations, with a view to better understanding the invasion history of mice in New Zealand, describing the spatial patterns of present genomic diversity, and the ancestral origins of each population.

## 3. Materials and Methods

### Sample collection

A total of 182 mouse tail samples ~10mm long were obtained from across the country (Figure 1), selected to achieve geographically representative sampling from across the two main islands, from all distant offshore islands with extant mouse populations, some large inshore islands, and from Sydney, Australia – a potential source population for invading mice, as it was the major port in the region in the 19^th^ Century. Where possible, samples of known mitochondrial lineages that had previously been sequenced for the mitochondrial control region by King et al., 2016 were used. Fifty-nine new samples were obtained from locations of interest that had previously not been sampled, or from locations where these previous tissue samples were found to be degraded. Tail samples were stored frozen from fresh.

**Figure 1.**
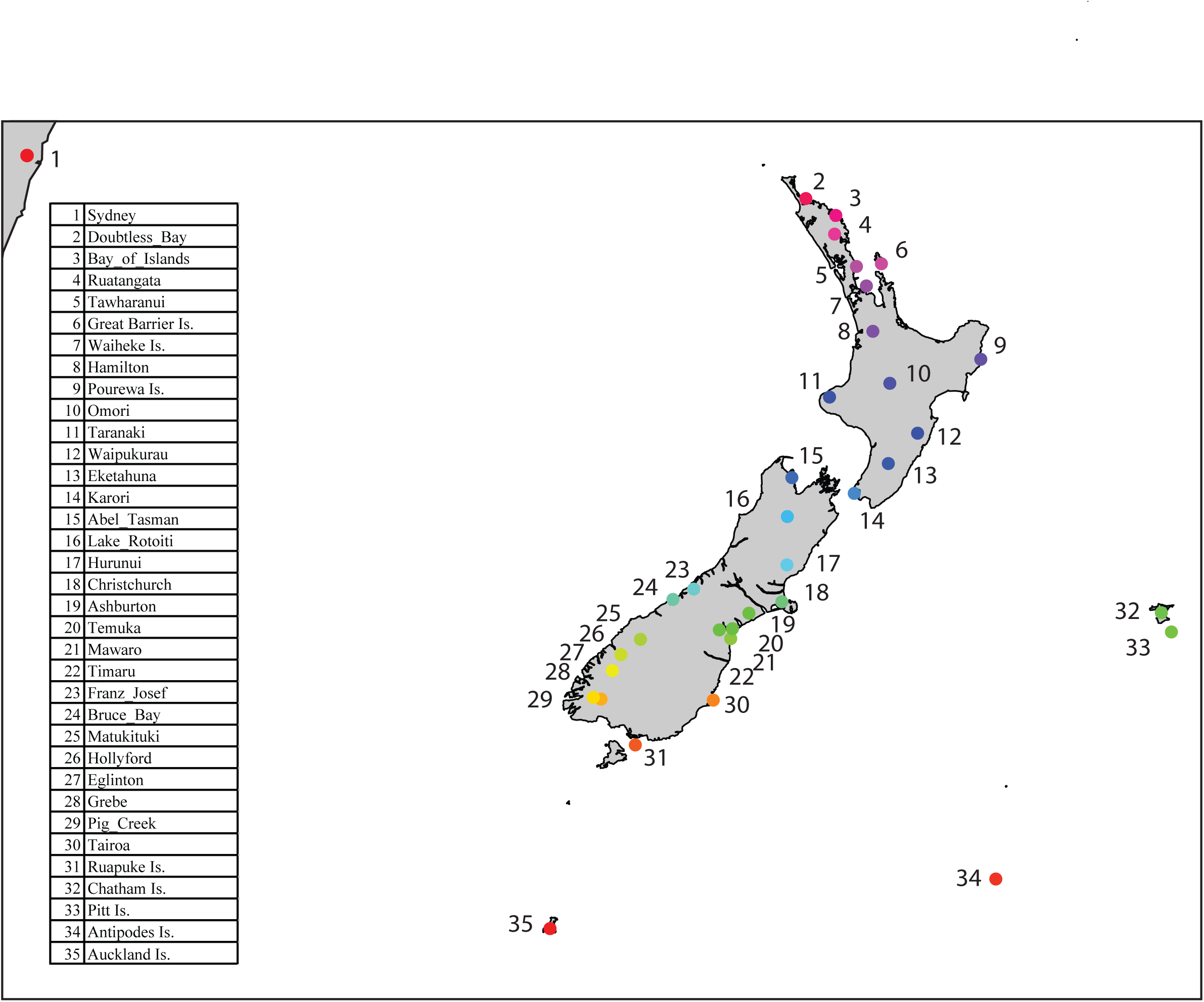
Sampling locations for genomic genotyping. For the mitochondrial dataset, data from King et al, 2016 was also included (q.v. for map and further details on those sample sites). Colour codes are based 011 ktitude and used to help display relationships between sampling locations in future figures.

### DNA extraction and Gigamuga sequencing

Tail samples were sent to the University of North Carolina, where genomic DNA was extracted using a Qiagen Gentra Pure tissue kit according to the manufacturer’s protocols. All genome-wide genotyping was performed using the GigaMUGA array at the University of North Carolina (GeneSeek, Lincoln, NE) [57]. Genotypes were called using Illumina BeadStudio (Illumina, Carlsbad, CA) and processed with ARGYLE [58].

### Bioinformatic filtering and analyses

We filtered and combined the separate genotyping runs in ARGYLE [58], an R package specifically designed for manipulating MUGA data. We then used PLINK 1.9 [59] to remove any individuals from the dataset that had 10% or more missing data, and to filter SNPs based on coverage across individuals (loci were retained only if present in at least 90% of samples). All further filtering and analyses steps were also conducted in PLINK 1.9 unless otherwise stated. We then merged our dataset with two published datasets: 1) the reference CigaMUCA dataset (with a number of loci identical to our data) and 2) the MegaMUGA wild mouse reference dataset. For the CigaMUCA dataset, we included only wild mice, and a few wild-derived lab strains that had been shown previously to be relatively pure and non-divergent from their sub-specific origins [57]. This dataset therefore retained only a handful of each subspecies as references, but contained the full set of ~135,000 SNP markers. The MegaMUGA SNP array consists of ~78,000 markers, of which ~65,000 overlap with the GigaMUGA array. Over 500 wild mice from across the native range have been genotyped using the MegaMUGA array, therefore this dataset provided a much larger wild reference dataset than the GigaMUGA this dataset provided a much larger wild reference dataset than theGreference dataset, but with reduced SNP coverage. Both our combined datasets were filtered for Linkage Disequilibrium (LD) with a window size of 10kb, a step size of 5 and an R^2^ of 2.

Both of these datasets were then filtered for a minimum minor allele frequency of 0.05. For subspecies identification, we further filtered the data to include only those loci which were most highly differentiated between subspecies (dataset 3). When the MegaMUGA array was developed, Morgan and colleagues (2016) evaluated the information content of each site in terms of subspecies differentiation – calculating the Shannon information content for each locus. This takes values between 0 and 1, where 0 means identical allele frequencies and therefore no information to inform identification, and 1 is reached when it detects a fixed difference between subspecies. We filtered our data for subspecies admixture calculations to exclude loci with a Shannon information of <0.5 – leaving only loci with high differentiation between subspecies, and removing the ascertainment bias in the chip towards *M. m. domesticus* diversity.

To investigate the population genetic structure within New Zealand, we used the program FASTSTRUCTURE [60], on the complete dataset (dataset 1) employing the choose.K command to ascertain the optimal number of clusters present in our data. To examine the autosomal (and X chromosomal) subspecies ancestry of individuals, we used the program ADMIXTURE, using both the combined MegaMUGA reference dataset (dataset 2), and the reduced, weighted dataset that contained only those SNPs that were most diagnostic for subspecies identification (dataset 3). For genome-wide comparisons in ADMIXTURE, we did some initial pilot runs using all of the reference samples, and then when it became clear that the majority of *M. m. domesticus* ancestry came from Europe, as expected from historical shipping records, we limited the subspecies reference dataset to include only wild *M. m. domesticus* from this region. This was done because working with highly uneven reference populations may cause biases in admixture assignment [61]. ADMIXTURE was run for all autosomal chromosomes together, and for each chromosome separately, to further resolve the contributions of each subspecies to the genomic makeup of each mouse. We then used the R package TessR3 to plot ancestry admixture coefficients spatially using Kriging [62].

As a comparison for the ADMIXTURE outputs, we also used the MegaMUGA wild mouse reference database to search for fixed differences (diagnostic SNPs) among subspecies reference sets, and then counted the relative contribution of these SNPs to each of the New Zealand mouse samples. While these diagnostic data yield a far smaller dataset than the total GigaMUGA or MegaMUGA genotypes, it provides an unbiased estimate of ancestral contribution, which can be compared to the model-based outputs from ADMIXTURE. We extracted both the mitochondrial and Y-chromosome SNPs and compared these haplotypes with the GigaMUGA reference samples, and with the known mitochondrial control region sequences previously recorded for most of the New Zealand samples [56]. There are multiple (>5) diagnostic SNPS on both the Y-chromosome and mt-genome featured on this array, therefore we were able accurately to classify each haplotype to subspecies origin, and where possible, to infra-subspecies clade.

We created an identity-by-state (IBS) differentiation matrix between individuals using PLINK, and used these to construct neighbour-joining trees in the R package APE [63], and principal component analyses (PCAs) in PLINK. We ran these analyses both for the New Zealand samples independently, and for the combined GigaMUGA (dataset 1) and MegaMUGA (dataset 2) references.

## 4. Results

### Data filtering and statistics

Of the 182 mouse tail samples collected from around New Zealand (and from Sydney and Lord Howe Island in Australia), 166 had high enough DNA quality to pass quality control (QC) and be analysed using the GigaMUGA SNP array. We filtered for a maximum of 10% missing data per individual, removing a further 5 individuals, yielding a final dataset of 161 mice. Neither of the two mouse samples obtained from Lord Howe Island were of high enough quality to be retained in analyses, but all other locations remained represented for spatial population analyses.

Of the 129,704 autosomal SNPs, 119,645 remained after filtering for coverage, and 49,266 remained for analyses requiring linkage equilibrium. Examining the reference samples, mitochondrial and Y-chromosome haplotypes could be assigned to subspecies using multiple (>5) fixed differences, and to intra-subspecies clade by >2 fixed SNPs. For mitochondrial haplotypes and clades for all individuals see Supplementary Table ST1. For the subspecies admixture analyses, we retained only SNPs that had a differentiation Shannon weighting of >0.5 between subspecies, leaving the most differentiated 9,501 SNPs for high-accuracy subspecies genomic assignment, and these were scattered across all chromosomes (Supplementary Table ST2).

As a second method to confirm ancestry proportions, we created datasets composed of fixed differences between subspecies pairs *domesticus*/*castaneus* (106 loci) and *domesticus*/*musculus* (481 loci). The relative number of these ‘fixed’ loci between subspecies will not reflect real differences in the levels of similarity between subspecies, nor are they necessarily fixed, because the sizes of the *M. m. castaneus* and *M. m. musculus* reference populations were small relative to those for *M. m. domesticus*. Given the large size of the *M. m. domesticus* reference dataset, these ‘fixed’ differences do however represent loci where one allele is likely to be very rare or absent from *M. m. domesticus*, therefore these should be useful for identifying ancestry from these other two subspecies.

### Genetic population structure

Across all sampling locations, individuals grouped together most closely with other individuals from the same location – see neighbour-joining trees (Supplementary Figures SF1, SF2), indicating that differentiation between locations was always higher than within them. There was also clear regional structure evident: for ease in describing the spatial genetic patterns of mice across New Zealand, we have divided the two main islands into five regions (A – E) corresponding to population genetic regions, and provide a map highlighting the locations mentioned in the text (Figure 2).

**Figure 2.**
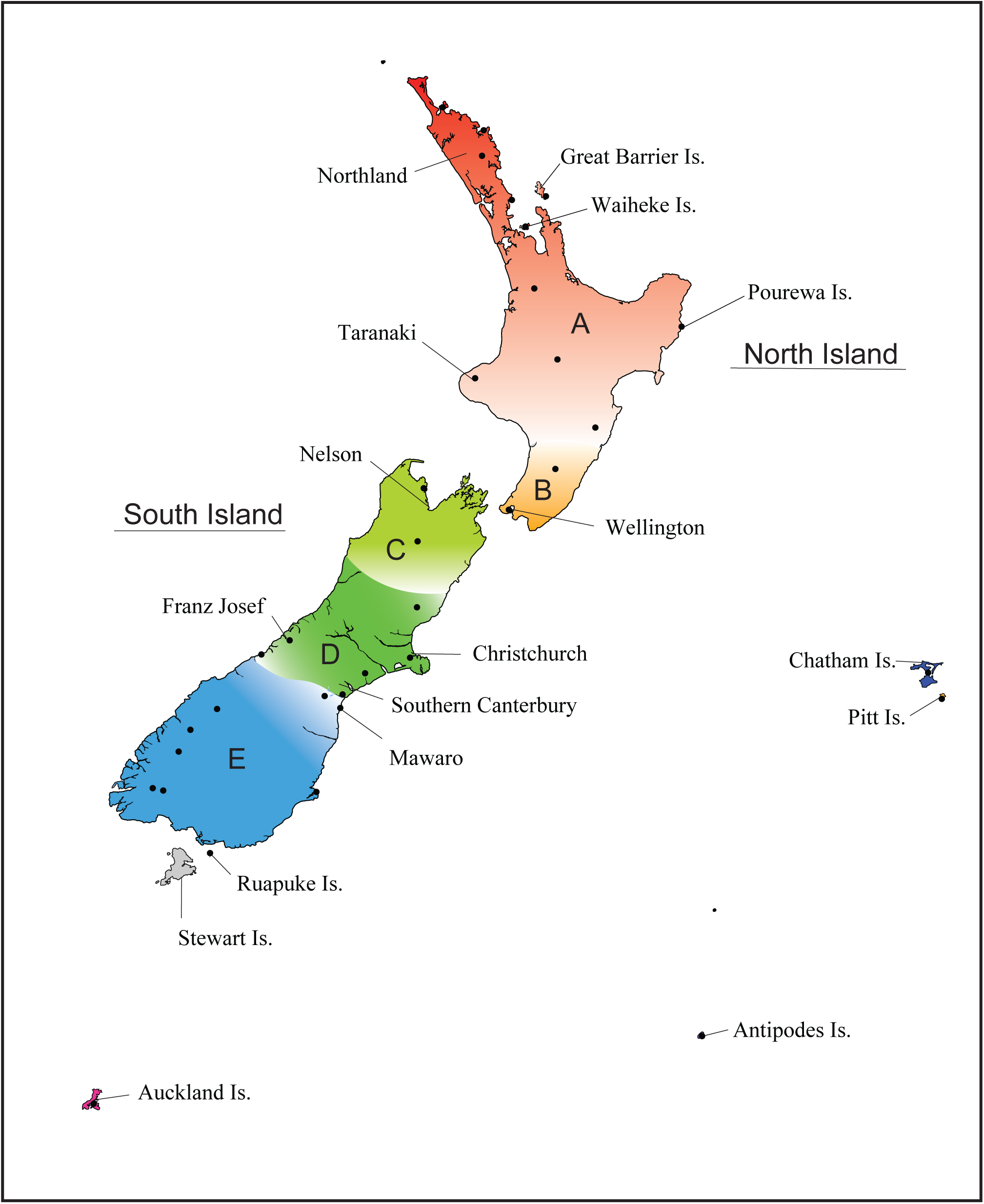
Map of New Zealand indicating the geographical regions discussed and highlighting any particular locations mentioned in the text. Sampling locations shown as black dots.

The primary population genetic structure among mouse populations in New Zealand is defined by the divergences between the southern South Island sampling locations (Matukituki, Hollyford, Eglinton, Grebe, Pig Creek and Tairoa - region E; for detailed location data see [56]), and the northern South Island sampling locations (Abel Tasman & Lake Rotoiti - region C) from the remaining locations (Figure 3). Sampling locations that exhibit admixtures with these divergent groups within the South Island (e.g. Hurunui, Bruce Bay), are shown as slightly divergent from the other populations.

**Figure 3.**
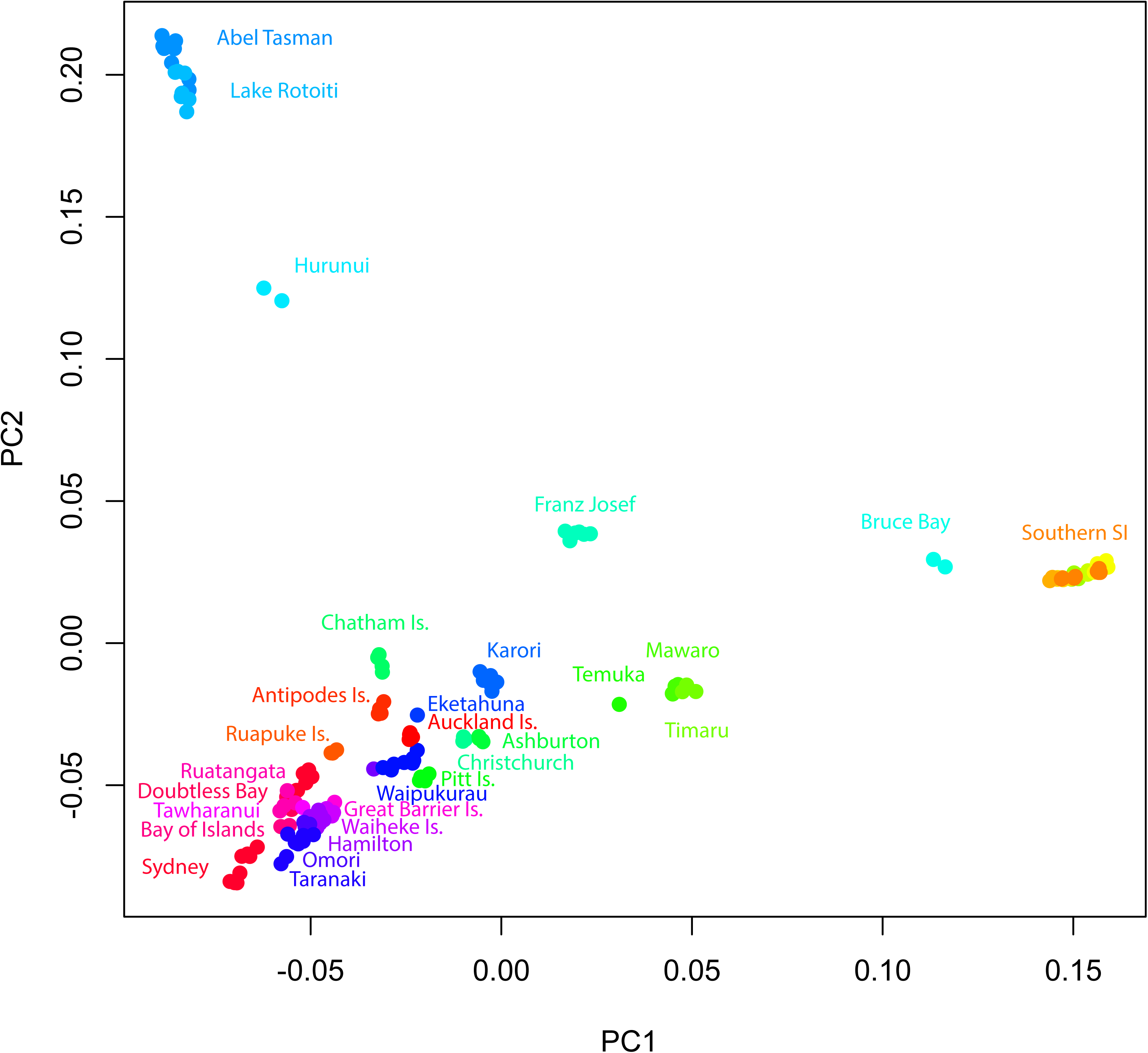
PCA based on IBD for all NZ mice derived from LD filtered loci. The southern South Island population (Southern SI) consists of Matukituki, Hollyford, Eglinton, Grebe, Pig Creek & Tairoa, which overlap too much to be labelled separately. Colours are derived from latitude, matching Figure 1.

While mice from each location could be identified to their sampling location (Supplementary Figures SF1, SF2), FASTSTRUCTURE indicated nine clusters were optimal to explain the genetic differentiation present across New Zealand (Figure 4A). These clusters represent groups of individuals with similar genetic makeup and similar ancestry – though there will be spatial patterns of diversity and connectivity within these groupings. It is possible that each cluster therefore represents a different population, founded primarily via different introduction events, though long periods of relative isolation could also account for the divergence of clusters.

**Figure 4.**
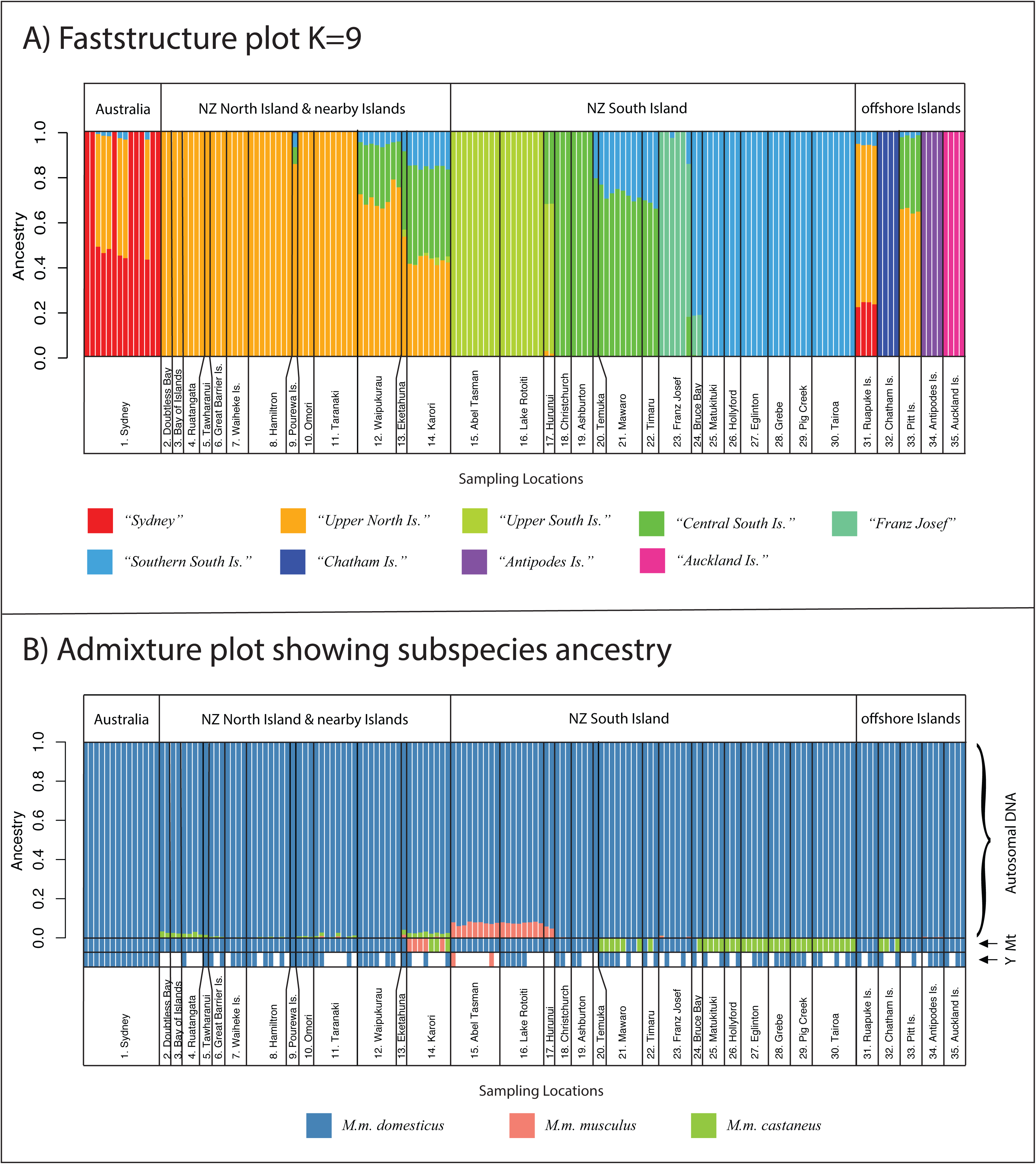
A) Cluster assignment plot derived from Faststructure (K=9) for New Zealand mice. Each individual mouse is represented by a column, and the proportion of each colour is the proportion of ancestry from that cluster. Clusters are named according to the region that primarily contributes to that cluster. B) Admixture plot showing the percentage of autosomal nuclear DNA ancestry from each subspecies for each mouse, along with their matrilineal mitochondrial (Mt) and patrilineal Y-chromosome (Y) ancestry. Each individual mouse is represented by a column, with the proportion of each colour representing the proportion of ancestry derived from each subspecies.

Three of the most remote offshore islands (Chatham, Antipodes and Auckland) are highly differentiated from all other populations. Ruapuke Island clusters with Sydney and some North Island sites, and Pitt Island is most similar to locations in the lower North Island. The relatively near-shore islands (Great Barrier, Waiheke and Pourewa islands) belong to the same cluster as nearby North Island mainland locations – all in region A.

For the mainland sites, admixture between clusters is evident, with southern Canterbury (Mawaro, Timaru and Temuka) being composed of a mixture of Central South Island cluster to the north (Christchurch, Ashburton – region D) and the southern South Island cluster to the south (Matukituki, Hollyford, Eglinton, Grebe, Pig Creek and Tairoa – region E). In the lower North Island – region B, a mixture of clusters is also evident, with contributions from the northern North Island cluster diminishing southwards, plus elements of both the central and southern South Island clusters.

### Genomic contributions from each subspecies

We found significant discrepancies among the mitochondrial, autosomal and Y-chromosome ancestries across the country, indicating frequent admixtures between subspecies and genetic clusters in multiple locations, both before and after arrival in New Zealand (Figure 4B).

Across all sampling locations, ADMIXTURE indicated that the nuclear ancestries of New Zealand (and Australian) mice are predominantly *M. m. domesticus*. In the southern North Island (region B), southern South Island (region E), and on Chatham Island, there were notable discrepancies between mitochondrial ancestry and nuclear ancestry (Figures 4B,5). In Wellington (Karori), all mice had either *M. m. musculus* or *M. m. castaneus* mtDNA, while their autosomal DNA consistently showed ~97% *M. m. domesticus* ancestry, with ~2% *M. m. castaneus* and ~0.02% *M. m. musculus* input. In the southern South Island *M. m. castaneus* mtDNA dominated, with all mice sampled south of Mawaro having *M. m. castaneus* mtDNA. ADMIXTURE however indicated no trace (<0.001%) of *M. m. castaneus* nuclear DNA in any individuals from these locations, and effectively no trace of *M. m. castaneus* nuclear ancestry across the South Island. This ADMIXTURE result matched closely the ‘diagnostic’ subspecies SNP frequencies, with 0.6% of ‘diagnostic’ *M. m. castaneus* alleles present on average across the southern South Island. Across all populations the diagnostic SNP marker sets confirmed the ADMIXTURE analyses (Supplementary Figure SF3), although since many of these loci are identical between *M. m. musculus* and *M. m. castaneus*, the proportions of hybrid alleles should be interpreted as a percentage of ancestry that is non-*domesticus* rather than clearly identifying one or other of these two minor component subspecies.

**Figure 5.**
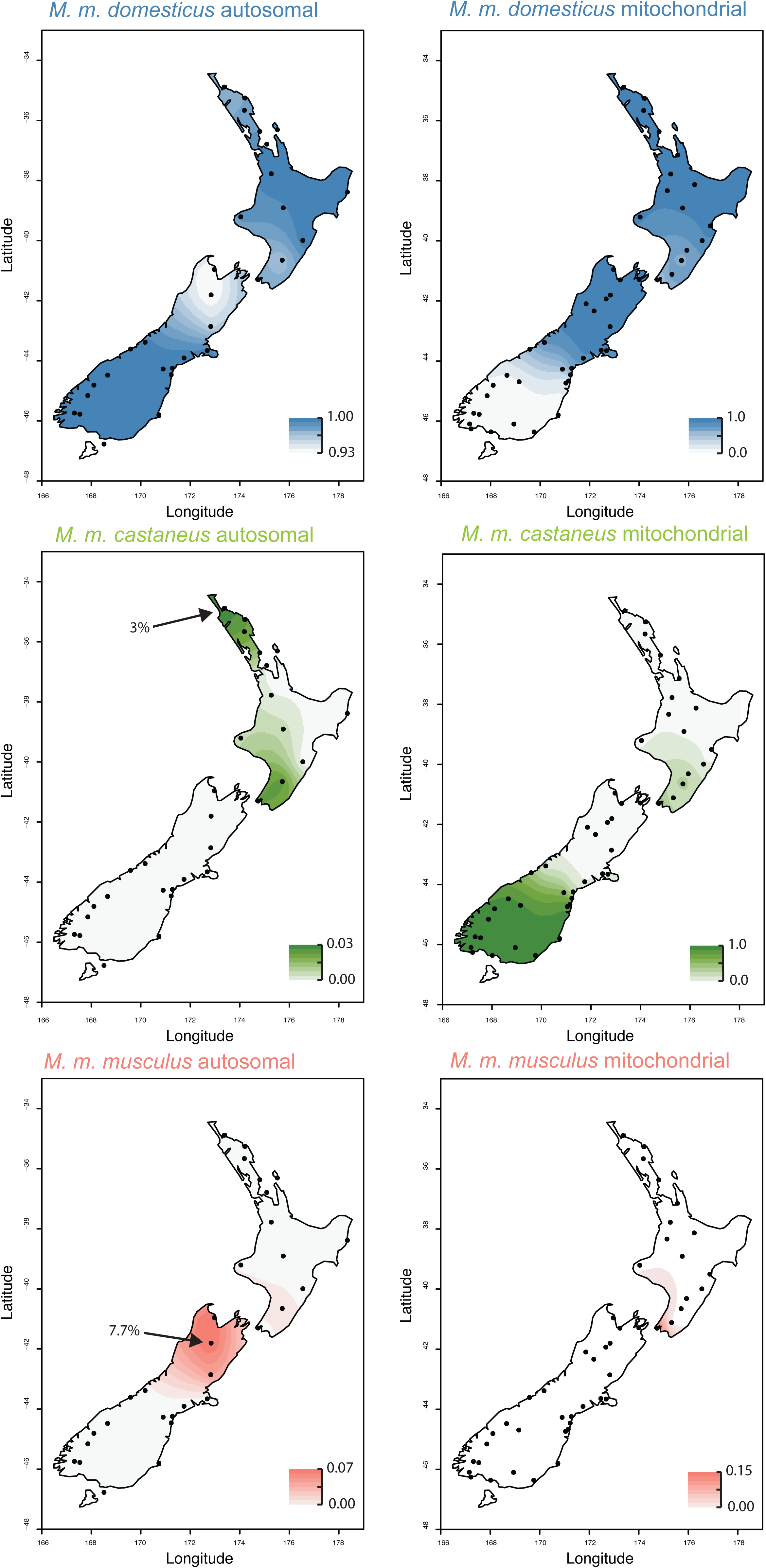
Maps of New Zealand showing the comparative subspecies ancestry proportions for mice in each region as determined in Tess3r for both nuclear autosomal DNA (left) and mitochondrial DNA (right). Note the different scales for each map, as ancestry percentages varied hugely between subspecies and DNA type. Mitochondrial data are concatenated from the present study and King *et al,* 2016. Sampling locations shown as dots, with the places with the maximum recorded nuclear ancestry for *M. m. castaneus* and *M. m. musculus* are indicated.

We detected a similar situation on Chatham Island, where three of the four mice sampled had *M. m. castaneus* mitochondrial DNA, and the fourth had *M. m. domesticus* mtDNA. No *M. m. domesticus* mtDNA had previously been recorded among 9 mice previously collected there. Nuclear ancestry of Chatham Island mice was consistently over 99.8% *M. m. domesticus* from the ADMIXTURE analysis. The diagnostic SNP analysis gave a slightly higher percentage of *M. m. castaneus* ancestry (~3%), though with the small number of loci available this may be less accurate than the ~9,500 SNPs analysed in ADMIXTURE. The *M. m. castaneus* mitochondrial genotypes from Chatham Island matched the previously identified casNZ.2 haplotype, and the single *M. m. domesticus* mitochondrial haplotype matched *M. m. domesticus* clade E haplotypes, which dominate both the North and South Islands.

The spatial distribution of subspecies ancestry, and the discrepancies between the mitochondrial haplotypes and nuclear genomes are highlighted in Figure 5 – note the differences in ancestry proportions in the scale bars between the comparative mitochondrial and nuclear maps. The only places in the country with any substantial *M. m. castaneus* contribution to the nuclear genome were in Northland (Doubtless Bay, Bay of Islands, Ruatangata, and Tawharanui – the northern part of region A), and the Wellington region (Karori and Eketahuna – region B), with 2-3% *M.m. castaneus* ancestry each (Figure 5). Traces (~1%) of *M. m. castaneus* ancestry were also recorded in Taranaki. Of these places, only Wellington had any evidence of *M. m. castaneus* mitochondrial DNA, and *M. m. castaneus* Y-chromosomal DNA was never recorded in any of the sampled locations.

While *M. m. musculus* mitochondrial DNA has never been recorded in the South Island, the three populations sampled in the north of the South Island (region C) showed a gradient of *M. m. musculus* autosomal ancestry from ~7-8% in Abel Tasman National Park and Lake Rotoiti, declining southwards to 5% at Hurunui. Two mice sampled from Franz Josef had ~1% *M. m. musculus* nuclear ancestry, suggesting that gene flow containing this admixed DNA has spread this far south. These observations were confirmed by the diagnostic SNP frequencies, and the *M. m. musculus* diagnostic alleles often clustered together on the genome, representing stretches of chromosomes inherited from this subspecies. The only two male mice sampled from Abel Tasman National Park both had *M. m. musculus* Y-chromosomes – and these were the only mice sampled across the entire study not to have *M. m. domesticus* Y-chromosomal ancestry.

Comparing the samples of mice from New Zealand and from the native range shows that all New Zealand mice clearly cluster with the wild native *M. m. domesticus* samples (Figure 6), though the populations with some *M. m. musculus* admixture (Lake Rotoiti, Abel Tasman, Hurunui – Region C) and *M. m. castaneus* admixture (Eketahuna, Doubtless Bay, Bay of Islands, Ruatangata, Tawharanui and Karori) are pulled slightly right, towards their respective minor subspecies components.

**Figure 6.**
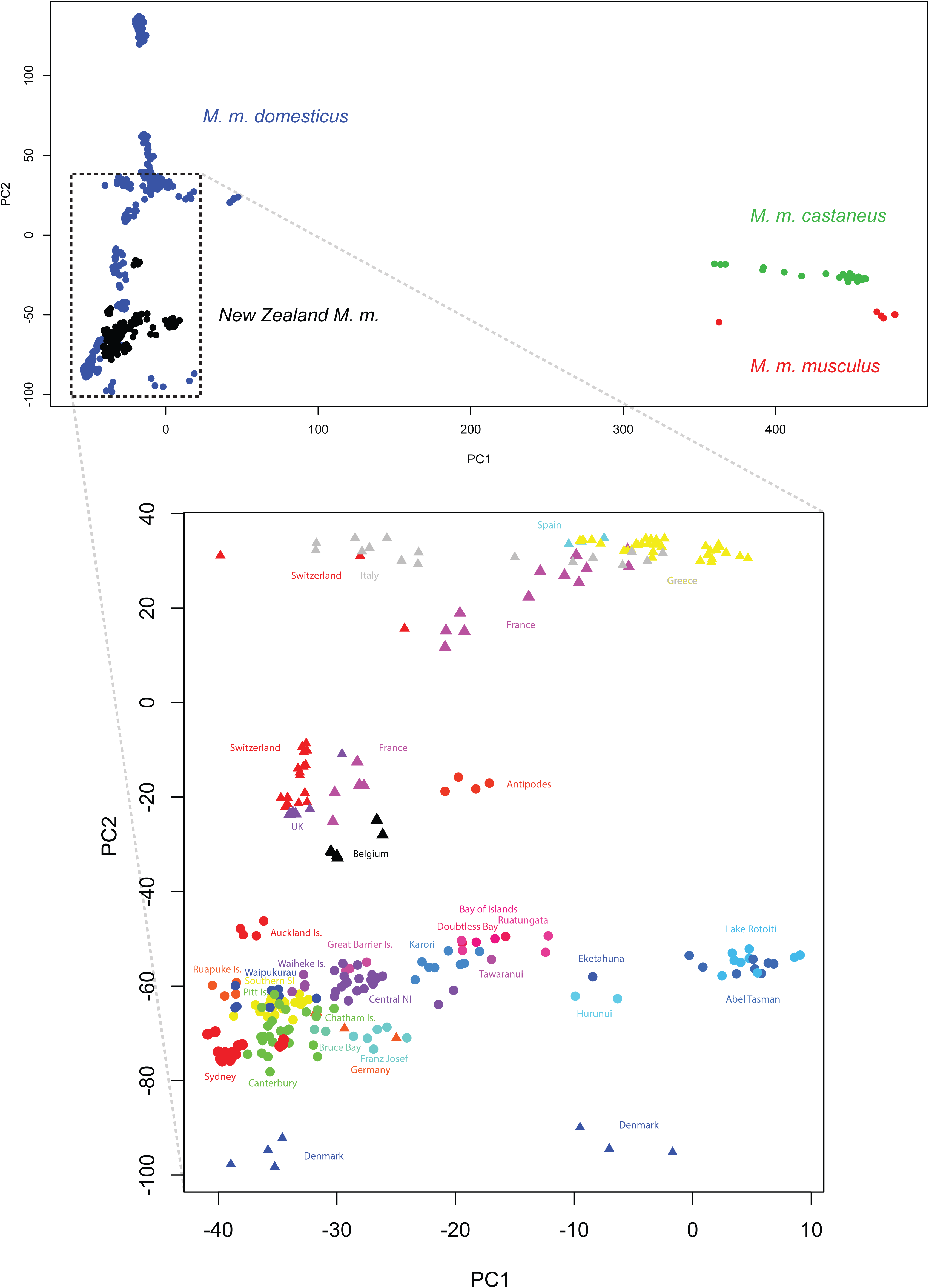
PCA based on sections of identity by descent for all wild mice genotyped with the MegaMUGA SNP array, with an enlargement below of the distribution of New Zealand wild mouse samples. New Zealand locations shown as circles, native-range samples shown as triangles (in the lower plot). Due to crowding we amalgamated some sampling locations for ease of display: Canterbury = (Christchurch, Ashburton, Mawaro, Temuka, Timaru), Central NI = (Hamilton, Omori, Taranaki, Pourewa Is.) and Southern SI = (Matukituki, Hollyford, Eglinton, Grebe, Pig Creek, Tairoa).

The contributions of each subspecies to the genomic makeup of New Zealand mice varied significantly across chromosomes (Figure 7). For the three identified geographic regions with large numbers of admixed individuals (at the nuclear level) – Northland, Wellington, and the upper South Island - we display these results graphically (Figure 7). Of particular note, there was minimal evidence for genomic input from *M. m. castaneus* or *M. m. musculus* for the X chromosome across these three populations, but a large proportion (>50%) of the genomic ancestry mice from the upper South Island came from *M. m. musculus* on chromosome 17, compared with the average *M. m. musculus* ancestry across the genome of around 7.5%.

**Figure 7.**
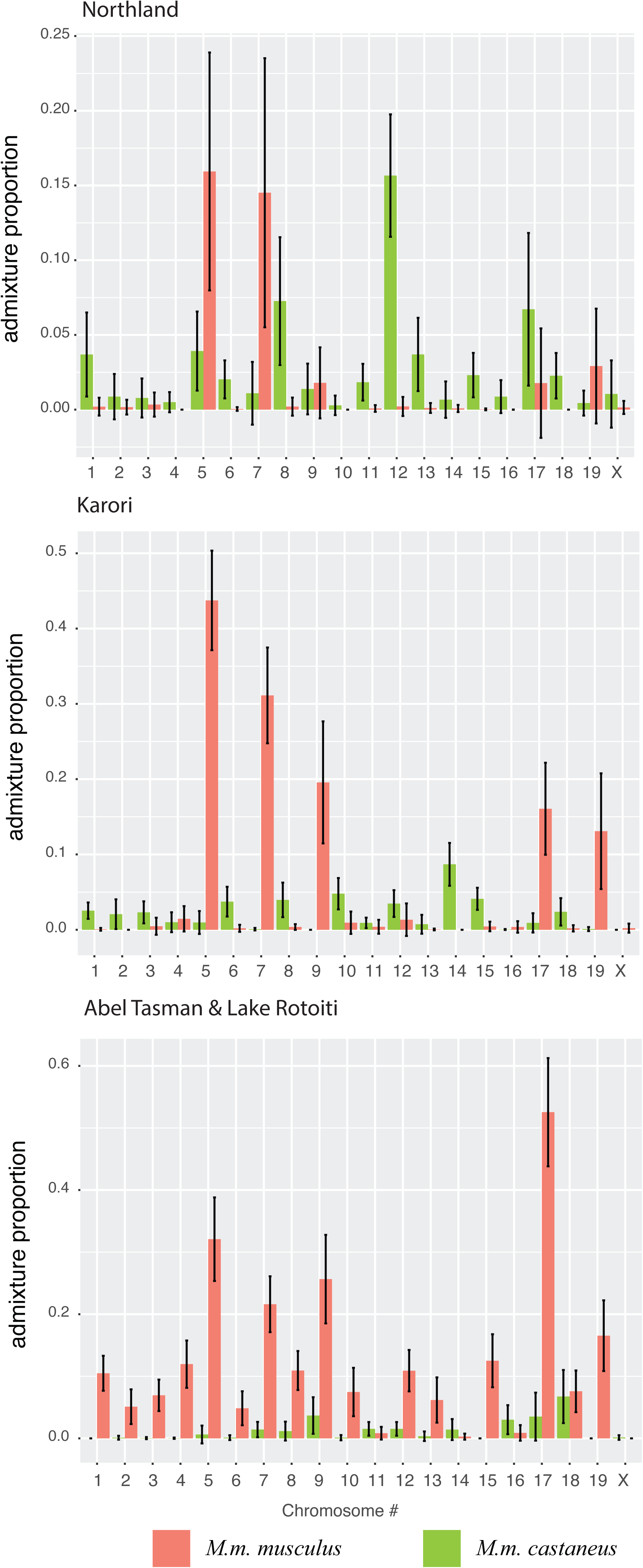
The proportion of admixture by chromosome for the three regions with highest nuclear diversity (Northland: Doubtless Bay, Bay of Islands, Ruatangata, Tawharanui; Karori; and the upper South Island: Abel Tasman & Lake Rotoiti). In all cases, only the admixture contributions of *M. m. castaneus* and M *m. musculus* are shown – the remainder being *M m. domesticus*.

## 5. Discussion

The application of cheap high-density genotyping arrays now available for mice has corrected many false assumptions, and greatly increased our knowledge about the diversity, ancestry and admixture of laboratory mice[46, 64]. These tools, developed primarily for developmental genetics and biomedical research, can also assist us in understanding the ancestry, invasion histories, and diversity of wild mice populations. Using the GigaMUGA SNP genotyping array, we have significantly expanded our knowledge of the genomic diversity and colonisation history of mice in New Zealand – highlighting the need to go beyond mitochondrial markers to trace biological invasions. This need for greater genomic resolution in evaluating biological invasions was recently also emphasized in a similar genomic study of Norway rats (*Rattus norvegicus*) [65] – another species for which invasion biology has benefitted from the genomic resources developed using domesticated laboratory strains. We have also gained significant insights into the abilities of wild mouse subspecies to hybridize during colonisation events.

### Insights into mouse subspecies hybridization

New Zealand is a particularly interesting location to look at the hybridization of mice sub-species in the wild, because traces of ancestry from all three major subspecies are present, the results of multiple comparatively recent hybridization events. The hybrid *domesticus*/*castaneus* populations in New Zealand are of particular interest given the rarity of this particular cross, however both of the previously identified ‘hybrid’ populations, one in the south of the South Island – region E, the other on Chatham Island, are hybrids only in the very limited sense that there is discordance between their nuclear *M. m. domesticus* and mitochondrial *M. m. castaneus* DNA. Both populations are essentially pure *M. m. domesticus* across the nuclear genome, but they retain the mitochondrial ‘ghosts’ of a previous hybridization event which failed to lead to nuclear admixture in the long term, i.e. they are cases of ‘mitochondrial lineage capture’ (reviewed in [66]. The fact that these populations have significantly different *M. m. castaneus* mitochondrial haplotypes implies that these mitochondrial lineage capture events occurred independently. The process of mitochondrial lineage capture has been observed across a diverse range of taxa including *Crotaphytus* lizards [67], chipmunks [68], loaches [69], deer [70], goats [71], hares [72-74], pocket gophers [75], voles [76], daphnia [77], and indeed between mouse subspecies [46, 78, 79]. While relatively commonly identified in mammals, mitochondrial capture with no traces of nuclear introgression has rarely been demonstrated to be so complete as in these mouse populations. Our ability to detect it has been made possible due to the extensive genomic markers available for this species.

Theoretical comparisons of pre- and post-zygotic models of isolation demonstrate that, under certain conditions, models of prezygotic isolation (e.g. female choice or male–male competition) allow for much more rapid introgression of maternally inherited DNA [80]. This result should be strongest when the source of the mtDNA is relatively rare, overall population sizes small, and there is asymmetric hybridization [66, 80]. These are precisely the conditions that would have occurred if a small founding population of resident *M. m. castaneus* was invaded by *M. m. domesticus* mice, and it is particularly likely given the relative hybrid fitness of these two subspecies.

The authors of reports of attempted crosses at the Jackson lab between *M. m. castaneus* and *M. m. domesticus* state that fighting is particularly prevalent in progeny of any crosses involving male *M. m. castaneus* [22]. Similarly, in the Collaborative Cross, a set of recombinant inbred lines derived from crosses between eight strains [81], the *M. m. castaneus* X chromosome was underrepresented [31]. Male infertility was responsible for nearly half of all observed lineage extinctions [82]. Furthermore, severe breeding problems have also been noted with crosses of another *M. m. castaneus* strain, CasA [83]. A recent quantitative trait loci (QTL) study of genes related to hybrid fitness identified regions on the autosomes, the X chromosome, and particularly in the Pseudo Autosomal Region (PAR) of the X and Y chromosomes which confer hybrid male sterility for crosses between *M. m. domesticus* and *M. m. castaneus* [31]. A substantial proportion of F_2_ males in White et al.’s study [31] exhibited phenotypes that previously had been connected with sterility. These included high levels of abnormal sperm, strong reductions in the apical sperm hook, and severely amorphous sperm heads that are unable to fertilize ova [41, 84, 85]. All of these factors indicate that when initially successful, hybridization between *M. m. domesticus* and *M. m. castaneus* is likely to be highly asymmetrical and unstable due to both behavioural and genetic incompatibility.

Along with the *domesticus*/*castaneus* hybrid populations described above, there is evidence of a hybrid population with nuclear introgression in the northern half of the South Island, between *M. m. domesticus* and *M. m. musculus*. The nuclear ancestry of this population is approximately 92% domesticus/8% musculus population, and Y-chromosomes from both subspecies are present in this region. These two Y-chromosomes could be spatially differentiated, because *M. m. musculus* Y-chromosomes were recorded only in the two male mice sampled from Abel Tasman National Park, while only domesticus Y-chromosomes were detected in five males from Lake Rotoiti. Given the small numbers sampled, we can only speculate about this trend. We have yet to find evidence of *M. m. musculus* mitochondrial ancestry in this population. Our results expand and quantify the findings of Searle et al. 2009, who also found some evidence of *domesticus*/*musculus* hybrids in this region.

Our finding that the *domesticus*/*musculus* hybrid population in the upper South Island has particularly high *M. m. musculus* ancestry for chromosome 17 is also an intriguing result worthy of further investigation. The only gene (Prdm9) known to cause hybrid sterility in vertebrates, identified in crosses between *M. m. musculus* and a classical inbred strain primarily derived from *M. m. domesticus*, is on chromosome 17 [27, 29, 86]. Our results could indicate that incompatibilities in this region have led to a high proportion of this chromosome being inherited from *M. m. musculus* across this population.

### The mouse invasion history of New Zealand

Our study highlights the extreme complexity of assessing the origins and invasion pathways of organisms using genetic data. While our results match those of previous studies [22, 49, 56] the vastly increased range of genetic markers highlight the need for genomic data to fully explain invasion histories. These previous studies relied on mitochondrial data, along with a handful of nuclear markers, because this focus allowed large numbers of mice to be genotyped. Since mitochondrial DNA is uni-parentally inherited, a clear and detailed pattern of inheritance can be established for this molecule and the matrilineal history [52]. Gene trees, however, are not species trees, and mitochondrial DNA is only one locus, which is largely unrelated to phenotype.

Firstly, we note the similarity of the nuclear genomes of mice across New Zealand. All mice in New Zealand other than those on Antipodes and Auckland Islands genetically clustered together with each other (and with Sydney, Australia), indicating similar origins, or significant mixing among locations post introduction. This pattern of similarity among most New Zealand sites (other than Antipodes and Auckland Islands) is more clearly seen in Supplementary Figures SF1, SF2. As previous work has indicated by the diversity of mitochondrial haplotypes, there have been many introductions to New Zealand of mice representing diverse origins, however, those that have contributed to the bulk of the modern nuclear genetic diversity across the country are primarily descended from *M. m. domesticus* ancestors from north western Europe.

Our study has revealed discordant genomes in many parts of New Zealand, as in other well-studied islands such as Madeira subject to multiple invasions by mice of different origin [87-94]. The differences between the genders in behaviour and breeding biology permit invading male markers to spread more rapidly than female markers [95]. Hence, island populations are more porous to incoming males than to females, so mtDNA is more likely to mark the original colonists. Therefore, mitochondrial DNA can be helpful in establishing priority among propagules in the order of colonisation, but misleading as to the genomic ancestry of individuals in the extant population. Here we update and review the story of the mouse invasion of New Zealand and its surrounding knowledge, in light of our new insights from the genomic data. King [55] made a number of hypotheses as to the origins of New Zealand mice, at that time based on mitochondrial data along with historical shipping records. We have reproduced a table of these hypotheses, along with the level of support offered by the genomic data (Supplementary Table ST3).

Briefly, the hypothesis of mice arriving to Sydney with supply fleets from Europe is highly supported, as all Sydney mice cluster with north western European mice – as had previously been suggested using mitochondrial data [50]. The hypothesis of mice arriving from India or Canton arriving to Sydney is not supported, as no *M. m. castaneus* nuclear or mitochondrial ancestry has been detected. If *M. m. castaneus* arrived in Sydney they either failed to establish, or were entirely replaced by *M. m. domesticus*. We cannot rule out traces of *M. m. castaneus* in small local populations around the ports - as our samples came from the north and west of Sydney, but if these exist, this genetic component must be minimal for traces to not have spread further.

The lack of *M. m. castaneus* signal in Sydney means that the *M. m. castaneus* ancestry recorded in New Zealand is likely to have come directly from Asia. There are clearly two *M. m. castaneus* mitochondrial lineages in New Zealand: 1) the southern South Island (region E) which is also present in the southern North Island (Region B), and 2) Chatham Island. Nuclear *M. m. castaneus* ancestry was however only recorded in the North Island, primarily in the southern and northern regions (Figure 5). There are several possible scenarios that may account for the first of these two hybrid populations (on the North and South Islands). King suggests 1) direct colonisation of the southern South Island from Canton by sealers in the 1790s to 1810s, 2) colonisation from trading ships from China to Wellington from 1840, and 3) from China to the southern South Island (Dunedin or Hokitika) with gold miners from 1865-1890 [55]. We cannot rule out any of these hypotheses, but given that the majority (or all of) the nuclear genome of mice in these regions has been replaced with *M. m*. domesticus DNA, hybridization must have started early, potentially before arrival on a boat or in a previous port.

The fact that it is the same mitochondrial lineage in the southern North Island and the southern South Island brings up the possibility that mice with *M. m. castaneus* ancestry colonised only one of these places, then moved to the other. This hypothesis is supported by the mixture of genetic clusters found in Wellington, including some ancestry for the southern South Island cluster. We have not yet tried to assess the direction of this movement.

In the northern North Island, the discordance between the same two subspecies runs in the opposite direction (Figure 4) with some traces of *M. m. castaneus* nuclear ancestry, but with no *M. m. castaneus* mitochondrial DNA yet detected. In the 107 mice from that area previously examined, 92% carried a single haplotype of *M.m. domesticus* identical with equivalent representatives of Clade E in UK and Australia. For compelling biological reasons summarised above, it is reasonable to doubt that *M.m. castaneus* could have invaded such a strongly established *M.m. domesticus* population in Northland. There are also historical reasons to suspect that mice arrived in the Bay of Islands only in the 1820s or 1830s, after restrictions on trans-Tasman trade with Sydney were lifted [55]. Sydney had by then developed into the major port of the southwest Pacific, offering unlimited opportunities for hybridisation among mice living on shore or among cargo. The most likely explanation for our results is that the mice colonising Northland and spreading south were already hybrids, dominantly *M.m. domesticus* but carrying evidence of past encounters with *M.m. castaneus*.

### Offshore Island mouse invasion histories

Mice from the three relatively nearshore islands off the north-east coast of the North Island (Great Barrier, Waiheke and Pourewa) all belonged to the same cluster as the nearby mainland, indicating probable colonisation from vessels moving between the mainland and each island.

Although Chatham and Pitt islands are relatively close to each other (~25 km) their mouse populations had different genetic histories. Pitt Island mice appear to be mixed from two clusters – the central South Island cluster, and the North Island cluster. Pitt Island mice had the mitochondrial D-loop haplotype DomNZ.7, and the only locations it has been found on the main islands of New Zealand are around Timaru – where the primary shipping company to Pitt Island is based. Our results therefore strengthen the view that mice may have been exchanged between these Pitt Island and the South Island [56]. Chatham Island mice, however, were very different. Mice from this population predominantly had an *M. m. castaneus* haplotype, casNZ.2, which has yet to be detected anywhere else in New Zealand or Australia. It remains difficult to speculate as to the origins of this population as there are no clear mitochondrial links, and nuclear clustering indicates it is very separate from other New Zealand populations. This differentiation may be due to high genetic drift and founding effects, or because these mice have (some) origins independent of the other New Zealand populations.

Three New Zealand southern island populations – Auckland, Antipodes and Ruapuke Islands supported mice belonging to clades different from the rest of the New Zealand mouse samples, indicating a probable origin outside of mainland New Zealand. Our genetic results lend strong support to two specific introduction scenarios for the Antipodes and Ruapuke mice.

All Antipodes Island mice so far sequenced were mitochondrial *M. m. domesticus* clade C, a clade originating from France, Spain, Portugal and Italy [19] that has yet to be detected on mainland New Zealand, or in Sydney – although a different Clade C haplotype is present on Ruapuke I*s*. The origins of the Antipodes mice appear to have an origin independent of the other mice in New Zealand, with strong inferred genetic links to France as the source of this invasion. As suggested by Russell [96] the Antipodes Island mouse population was probably founded through a shipwreck, and a likely contender is that of the the *Président Felix Fauré*, a four-masted barque which was wrecked on rocks on the north side of the island in Anchorage Bay in 1908. All 22 men on board made it ashore and survived for two months before being rescued [97, 98]. The first records of mice on Antipodes Island are dated to one year later in 1909, by Waite [99] who wrote, “I am told by Captain Bollons that mice are very numerous at the Government depots on Campbell and Antipodes Islands”.

The genomic links between the Ruapuke Island population and mice from Sydney also match the known invasion history of the island. The first recorded population of mice in New Zealand arrived on Ruapuke in 1824 with the stranded flax trading ship *Elizabeth Henrietta*, which came from Sydney [100]. The fact that they have a mitochondrial haplotype of Clade C not yet observed in Sydney (or mainland NZ) could be due to (1) the small number of samples available of mitochondrial haplotypes from Australia, which are few and not from around the historical dock area; or (2), founding effects whereby a small random sample of a relatively rare haplotype in Sydney rose to prominence on Ruapuke Island.

The origin of the Auckland Island mice remains less certain. Following the discovery of the Auckland Islands in 1806, mice were first recorded there in 1840 by a United States expedition, but likely had already been present for some time before this. As there are no records of shipwrecks during this period, it is speculated that mice arrived here during sealing activities [101]. The only mitochondrial haplotype found on Auckland Island (NZ_dom4) is from clade E and matches haplotypes from Sydney, and both North and South Islands of mainland New Zealand. At a nuclear level, however this population clades most closely with introduced mouse populations from the USA. This population was possibly founded through activities of American sealers (or whalers) which were both active in the region at the time, although, due to the very significant bottleneck experienced by the Auckland Island population, further research and modelling will be needed to reveal the source of the mouse population on Auckland Island.

### Applicability of the Gigamuga SNP array

Ideally for population genomic studies, SNP variation recorded should represent the average SNP variation present across individuals, however this is rarely the case. SNP genotyping often suffers from an ascertainment bias, due to the procedure used to select SNPs [102-107]. The degree of ascertainment bias primarily depends on the size and representativeness of the panel of individuals, in this case the mice, used to select the SNPs. If a panel is chosen from individuals from a subpopulation or geographic region that is not representative of the population as a whole, variability in groups related to the ascertainment group will be over-represented [106].

The ascertainment bias for the Gigamuga SNP array is both extreme and not uniform, due to the development procedures used to create the array. The Gigamuga array was designed primarily from lab strains of mice, for use in biomedical and developmental genomic research. As the majority of laboratory strains are descended primarily from *M. m. domesticus*, a large proportion of SNPs will therefore be informative only between *M. m. domesticus* lineages, and monomorphic in the other subspecies. Furthermore, SNPs in the array were selected in a way to minimize mutual information between markers, which has the side effect of eliminating linkage disequilibrium (LD) signals. This ascertainment bias means that comparing populations with differing proportions of the three subspecies may lead to biases in estimates of nucleotide diversity, population size, demographic changes, linkage disequilibrium, selective sweeps and inferences of population structure [108-110]. We specifically chose analyses and data-filtering steps to avoid the effects of the inherent ascertainment bias in the GigaMUGA array, and these methods should be robust to the previously mentioned caveats [107, 110]. We caution against using our data to assess other properties such as nucleotide diversity or to identify selective sweeps without careful consideration for appropriate filtering.

### Future directions

Our study is the first looking at the invasion history of wild mice using the GigaMUGA SNP array, and indeed the first to use a high-density SNP array of any kind to assess population genomics of wild mice. Our data are available online (dryad.tm617) so that researchers, both from ecological and genomic fields can compare their populations with ours using similar SNP genotyping methods. For regions within the native range of house mice where complex patterns of divergence and introgression have been observed, such as Turkey and Iran [111, 112] and across Europe [20], the GigaMUGA array has significant potential to add to our understandings of the genomics of these hybrid zones.

We have not fully addressed the precise origins in the native ranges of the representatives of each subspecies that came to New Zealand, largely due to the paucity and unevenness of wild-mouse samples genotyped across these native ranges. However, these results are consistent with what is known from historic shipping records [55]. Given the standardization of the GigaMUGA array, it should be relatively easy to investigate this in the future, by obtaining and genotyping samples of mice from across the home range – particularly around historically significant ports. Furthermore, using runs of homozygosity, it should be possible to model the demographic history and the effects of the expanding invasion fronts on genomic diversity (e.g. [113]).

New Zealand is currently investigating the use of gene-drive technology [114] to help eradicate mammalian pest species as part of the aspirational “predator free 2050” project [115]. Current laboratory proof-of-concept research is proceeding on mice as a model organism, due to their short generation times and the extensive genomic resources available for this species. Our study of wild mice would be informative to this research, as an understanding of the standing variation, spatial structuring and genomic ancestry of wild mice will be vital to informing laboratory work, and could help identify islands where trials or application for this technique could be conducted.

## Acknowledgments

Thanks to all the people who supplied mouse specimens for the previous mtDNA analyses, and to Neil Barton, Anna Clark, Daniel Hurley, Pete McClelland, Peter Banks, John Birchfield, Jean Stanley, Russel and Teresa Trow, Judy Gilbert, Lyndon Slater, Elaine Murphy, John Henderson, Jamie Mackay for newly collected material. Thanks also to Anne Mitchell for sample curation, and to Andrew Morgan, Darla Miller at UNC for their extensive support in the sequencing, filtering and advice for analyses. Funding for genetic analyses supplied by the University of Waikato Strategic Investment Fund Grant F715 to Carolyn King.

## Ethical Statement

Tissue samples obtained for this research were either from pre-existing collections, or were obtained from pest control contractors.

## Data Accessibility

The software used in the research can be located as follows:

Plink 1.9: https://www.cog-genomics.org/plink2

Argyle: https://github.com/andrewparkermorgan/argyle

Admixture: https://www.genetics.ucla.edu/software/admixture/download.html

Tess3r: https://github.com/cayek/TESS3/tree/master/tess3r

Ape: https://cran.r-project.org/web/packages/ape/index.html

The reference MUGA data in PLINK format can be obtained at: http://datadrvad.orq/resource/doi:10.5061/drvad.689d2.

All raw genotypic data described in the manuscript have been uploaded as PLINK binary files (.bed, .bim, .fam) to Dryad.

## Authors’ Contributions

Andrew Veale resampled all the accumulated material stored at Waikato University described by King et al 2016; extracted and processed all new genetic samples; sent them for processing; analysed the results; and wrote most of the paper.

James Russell supplied extra samples from Abel Tasman, Antipodes Is., Auckland Is., Ruapuke Is., Tawharanui, Bay of Islands, Doubtless Bay, Timaru, Pourewa Is. and Lord Howe Is., and commented on the MS.

Carolyn King organised the collection and care of the mice since 1999; applied for the funds; and commented on the MS.

